# Combinatorial regulation of cellular rotation during dorsal-ventral axis establishment in *Caenorhabditis elegans*

**DOI:** 10.64898/2026.06.28.735104

**Authors:** Mi Jing Khor, Caleb Lai, Chelsey Lynn Gough, Yuxuan Rain Xiong, Aoi Hiroyasu, Taile Li, Viktorija Juciute, Christina Rou Hsu, MinJee Kim, Kalen Dofher, Kenji Sugioka

**Affiliations:** Life Sciences Institute, The University of British Columbia, 2350 Health Sciences Mall, Vancouver, BC V6T1Z3, Canada; Department of Zoology, The University of British Columbia, 2350 Health Sciences Mall, Vancouver, BC V6T1Z3, Canada

## Abstract

Cellular rotation is an understudied mechanism that regulates animal morphogenesis. In *C. elegans*, the dorsal-ventral axis is established when the two-cell-stage AB cell rotates within the eggshell as it divides, generating the diamond-shaped blastomere arrangement at the four-cell stage. Multiple mechanisms, including actomyosin-dependent oriented division, chiral cortical flow, and eggshell shape, have been proposed to regulate this arrangement, but whether these represent conflicting hypotheses or co-acting mechanisms remains unclear. Here, we show that CUL-3-actomyosin-dependent oriented division, eggshell geometry, and the Ras-MAPK signaling pathway regulate distinct steps of cellular rotation. AB cell rotation occurs in two distinct phases: Phase I during AB cytokinesis and Phase II during neighboring P_1_ cytokinesis. We found that CUL-3-actomyosin-dependent oriented division, eggshell geometry, and Ras-MAPK signaling are required for Phase I rotation. Errors in Phase I can be fully corrected by subsequent Phase II rotation. However, perturbation of CUL-3-actomyosin-dependent oriented division compromises this robust fail-safe mechanism, leading to dorsal-ventral axis defects. These results show the combinatorial and robust regulation of cellular rotation and its critical role in *C. elegans* embryogenesis.

## Introduction

Cellular rotation is an emerging morphogenetic mechanism by which cell-intrinsic polarity, cytoskeletal chirality, cell-cell adhesion, external geometry and developmental signaling are translated into stereotyped tissue architecture and body-axis formation (Asan et al., 2016; Naganathan et al., 2014; Pohl and Bao, 2010; Wang et al., 2013).

A prominent manifestation of cellular rotation in morphogenesis is dorsal-ventral axis (D-V axis) establishment in *C. elegans* during the second and third mitotic divisions (Rose and Gönczy, 2014). At the 2-cell stage, the mitotic spindle of the anterior blastomere, AB, is oriented orthogonally to the anterior–posterior axis (A-P-orthogonal axis), whereas that of the P_1_ cell is oriented along the anterior–posterior axis (A-P axis) (Figure 1A). As the AB cell enters anaphase and begins to elongate, it undergoes rotation within the eggshell. When the embryo is compressed, the entire embryo initially undergoes axial rotation around the A-P axis, a process outside the scope of this study. Subsequently, regardless of compression, the AB cell itself rotates, causing its division axis to deviate from the A-P-orthogonal axis. The rotational AB division drives displacement of the cytokinetic midbody from the first mitosis toward the future ventral side (Singh and Pohl, 2014). This midbody can then orient the P_1_ mitotic spindle, leading to an anteroventrally oriented P_1_ division. After completion of AB division and subsequent P_1_ division, the posterior daughter of AB (ABp) and the anterior daughter of P_1_ (EMS) align perpendicular to the A-P axis (Figure 1A). The resulting four-cell-stage arrangement is diamond-shaped, with the ABa and P2 cells not forming cell–cell contact. Ultimately, contact with P_2_ activates Wnt signaling in EMS and Notch signaling in ABp, resulting in D-V axis establishment and distinct cell fate specification (Rose and Gönczy, 2014).

**Figure 1.**
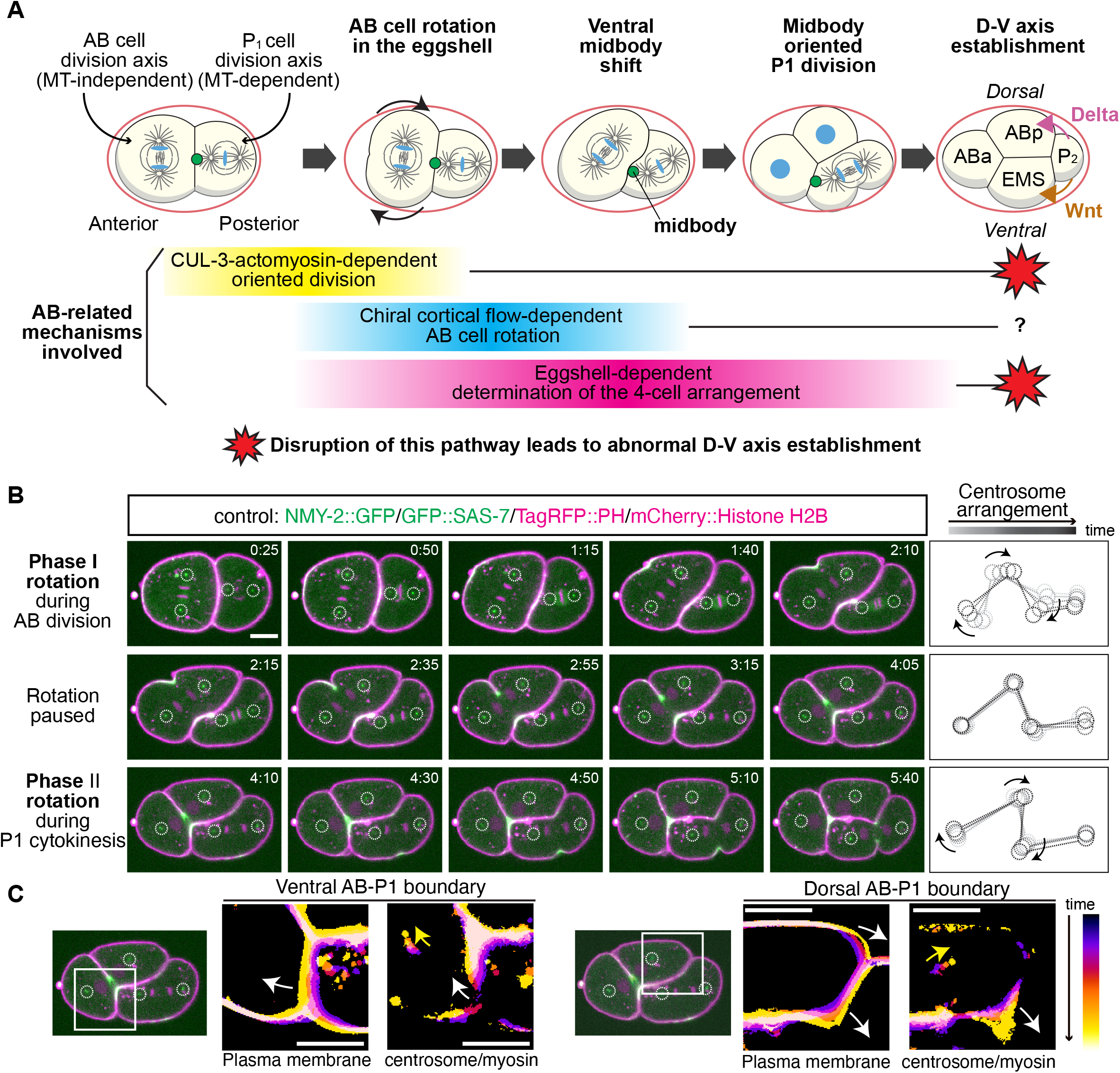
Dorsal-ventral axis establishment involves two phases of cellular rotation in *C. elegans*. **(A**) *C. elegans* dorsal-ventral axis establishment and the mechanisms implicated in AB cell rotation (see text for details). (**B**) Two phases of cellular rotation during AB and P_1_ divisions. NMY-2 (green, myosin), SAS-7 (green, centriole), TagRFP::PH (magenta, plasma membrane), mCherry::Histone H2B (magenta, chromosome). Dotted circles indicate centrosome positions. Right panels show the trajectories of each centrosome and lines connecting key centrosomes. The line connecting the posterior AB and anterior P_1_ centrosome serves as an indicator of the dorsal-ventral axis. Times are minutes and seconds relative to anaphase onset. (**C**) Temporal color-coded images of the plasma membrane, centrosomes, and boundary myosin at the ventral and dorsal AB-P_1_ boundaries. White and yellow arrows indicate the boundary and centrosome trajectory, respectively. Scale bars, 10 µm.

The orthogonal division axes patterns of AB and P_1_ are regulated by microtubule-independent and microtubule-dependent mechanisms, respectively. The A-P axis-oriented P_1_ division depends on the microtubule-pulling force generators LIN-5/NuMA and GPR-1/2 (Lorson et al., 2000; Srinivasan et al., 2003). In contrast, the A-P-orthogonal axis-oriented AB cell division requires the CUL-3 E3 ubiquitin ligase and actomyosin (Sugioka and Bowerman, 2018). The correct AB division axis is instructed by cell contact with P_1_, which is not altered by microtubule depolymerization. Furthermore, attachment of adhesive bead to the isolated AB cell orients its division axis toward the cell-bead contact site in an actomyosin-dependent manner, producing the cell arrangement similar to the A-P-orthogonal division axis (Sugioka and Bowerman, 2018). Hereafter, this cell contact and actomyosin-dependent mechanisms is referred to as CUL-3-actomyosin-dependent oriented division. Cortical microtubule-pulling force generator LIN-5 is also expressed in the AB cell and contributes to AB centrosome separation in prophase and compression-dependent AB cell rotation around the A-P axis in early cytokinesis (Bondaz et al., 2019; Middelkoop et al., 2024). However, LIN-5 is likely not involved in establishing the A-P-orthogonal division axis at anaphase onset, as LIN-5 depletion or low-concentration Nocodazole treatment does not affect the AB division axis, unlike the P_1_ division axis (Sugioka and Bowerman, 2018).

Inhibition of CUL-3–actomyosin-dependent oriented division results in defects in the AB division axis and dorsal-ventral axis establishment (Sugioka and Bowerman, 2018). After close inspection of live-imaging datasets, we suspected that the dorsal-ventral axis establishment defects arise from impaired AB cell rotation. Among the numerous mechanisms and phenomena reported at this embryonic stage, we identified two additional mechanisms that may contribute equally, or even more substantially, to AB cell rotation. The first mechanism is regulated by twisting behavior of the cell cortex movement, termed chiral cortical flow. When the AB cell divides, two hemispheres of the daughter cell surface counter-rotate around the spindle axis and the degree of counter-rotation is described as chiral cortical flow velocity. Reduction or enhancement of chiral cortical flow velocity correlates with reduced or increased AB cell rotation, respectively (Pimpale et al., 2020). Chiral cortical flow is limited to the AB cell and its descendants and does not function in P_1_ lineage (Pimpale et al., 2020). A second mechanism is eggshell-dependent. For decades, it has been known that the eggshell and vitelline membrane are required for the correct 4-cell-stage arrangement (Schierenberg and Junkersdorf, 1992). Recent studies further show that the eggshell geometry controls the 4-cell-stage arrangement (Seirin-Lee et al., 2022; Yamamoto and Kimura, 2017). For both chiral cortical flow and eggshell-dependent mechanisms, their requirements for A-P-orthogonal division axis in the AB cell remain unexplored. Furthermore, whether these three mechanisms represent conflicting hypotheses or coordinately acting mechanisms remains unclear. More specifically, it remains possible that the CUL-3–actomyosin-dependent oriented division mechanism we previously reported may not be involved in AB cell rotation, unlike other proposed mechanisms. Therefore, determining which pathways are required for division-axis orientation and cellular rotation is essential for further understanding dorsal-ventral axis establishment in this important model system.

In this study, we quantitatively analyzed AB cell behavior during cell division using live imaging. We identified distinct phases of AB cell rotation (Phase I and Phase II) and analyzed the contributions of the aforementioned pathways to each step, as well as to A-P-orthogonal division axis before the onset of rotation. We further examined the role of CUL-3-actomyosin-dependent oriented division using adhesive beads and clarified its seemingly separable roles before and during AB cytokinesis. Finally, we investigated the requirement for Ras–MAPK signaling, a novel regulator of AB cell division discovered in our recent RNAi screen, in this process.

## Results

### *C. elegans* dorsal-ventral axis establishment involves two phases of cellular rotation

To investigate the mechanism of D-V axis establishment, we precisely characterized the cellular rotation process using live imaging. Throughout the manuscript, we use the term “rotation” specifically to refer to cellular rotation, distinguishing it from spindle orientation. After anaphase onset, the AB cell began to elongate and initiate rotation within the eggshell, as previously known (Figure 1B; top row). This initial rotation lasted for approximately 1 min (65.8 ± 18.2 s, n = 20); however, we found that the AB cell then paused for approximately 1.5 min (92.1 ± 21.5 s, n = 20) before resuming rotation when the P_1_ cell underwent cytokinesis. We therefore designated these two distinct steps of AB cell rotation as Phase I rotation and Phase II rotation, respectively (Figure 1B). During Phase II rotation, the ventral AB–P_1_ boundary moved anteriorly, whereas the dorsal AB–P_1_ boundary moved posteriorly, coinciding with the corresponding movements of the AB centrosomes (Figure 1C). This observation may indicate that P_1_ cytokinesis rotates the AB cell by pushing the ventral AB–P_1_ boundary and pulling the dorsal AB–P_1_ boundary. To test the requirement of P_1_ cytokinesis in Phase II rotation, we delayed P_1_ division by RNAi knockdown of DIV-1, the homolog of the B subunit of the DNA polymerase α-primase complex (Fig. S1A), whose knockdown significantly delays P_1_ division (Fig. S1B) (Encalada et al., 2000). We found that Phase I AB cell rotation occurred normally while P_1_ cells remained in prophase throughout this process (Figures S1A, top row, and S1C). This rotation was followed by a small reversal as the P_1_ cell shrank during mitotic rounding, further supporting the influence of P_1_ cell shape on AB cell rotation (Figure S1A, second row from top). After a pause, Phase II rotation occurred to a similar extent as in wild type (Figures S1A, bottom row, and S1D). However, the timing of Phase II rotation was significantly delayed in *div-1(RNAi)* embryos. Consistent with a role for P_1_ cytokinesis, the onset of P_1_ cleavage furrowing and Phase II rotation were strongly positively correlated (Pearson’s r = 0.97, p < 0.0001, n = 30). These observations suggest that *C. elegans* dorsal–ventral axis establishment involves two phases of AB cell rotation, the second of which depends on P_1_ cytokinesis.

### CUL-3-actomyosin-dependent oriented division controls the initial AB division axis

Previous studies have suggested three mechanisms that contribute to AB cell rotation: CUL-3–actomyosin-dependent oriented division, chiral cortical flow, and an eggshell-shape dependent mechanism. However, their relative contributions to Phase I and Phase II rotation, as well as to establishment of the A–P-axis-orthogonal division axis before the onset of rotation, have not been comparatively analyzed (Figure 2A).

**Figure 2.**
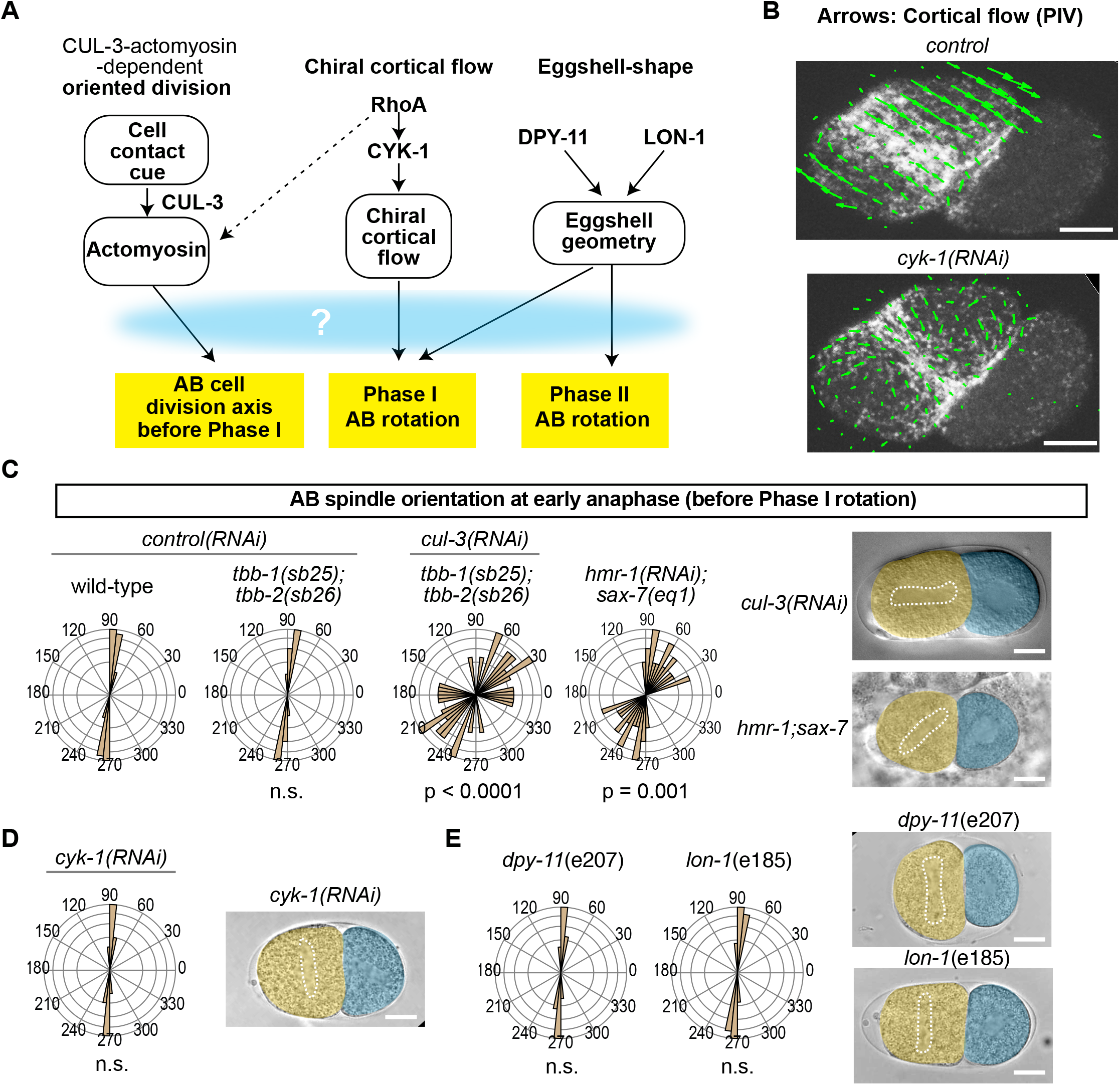
The initial AB division axis is regulated by CUL-3–actomyosin-dependent oriented division but does not depend on chiral cortical flow or eggshell geometry. (A) Three mechanisms implicated in AB cell rotation. How each pathway contributes to distinct aspects of AB cell dynamics remains unclear. Arrows indicate inferred contributions based on previous literature. (B) CYK-1-dependent cortical flow during AB cytokinesis. Myosin::GFP movement was analyzed using particle image velocimetry. Arrows indicate the direction of cortical flow. In control embryos, cortical flow was oriented in opposite directions on the ventral and dorsal sides of the daughter cell, resulting in counter-rotation. This counter-rotating flow was lost in *cyk-1*(RNAi) embryos. (C–E) AB spindle orientation in early anaphase before Phase I rotation following inhibition of the CUL-3–actomyosin-dependent oriented division pathway (C), inhibition of chiral cortical flow (D), or manipulation of eggshell geometry (E). *P* values were calculated using the Mardia–Watson–Wheeler test for equal distributions. Scale bars, 10 µm.

We first analyzed the roles of these mechanisms in A-P-orthogonal division axis orientation during early anaphase, immediately before Phase I rotation. To perturb CUL-3-actomyosin-dependent oriented division, we knocked down the CUL-3 E3 ubiquitin ligase, which was previously shown to be involved in contact-dependent regulation of actomyosin dynamics in the AB cell (Sugioka and Bowerman, 2018). As *cul-3(RNAi)* in a wild-type genetic background causes small mitotic spindles due to defective degradation of the microtubule severing protein MEI-1/katanin, we suppressed this phenotype using the suppressor tubulin mutant background *tbb-1(sb25);tbb-2(sb26)* (Lu and Mains, 2005). *tbb-1(sb25); tbb-2(sb26)* mutants alone exhibited no detectable phenotype (Figure 2B). In contrast, *cul-3(RNAi)* in this background caused defective AB spindle orientation, as we reported before (Sugioka and Bowerman, 2018). Hereafter, all *cul-3(RNAi)* experiments were performed in the suppressor tubulin mutant background. To provide further evidence that the AB division axis is regulated by cell-cell contact, we depleted the adhesion proteins classical cadherin HMR-1 and SAX-7/L1CAM, which are major early embryonic adhesion proteins (Grana et al., 2010). In *hmr-1(RNAi); sax-7(eq1)*, the AB division axis was also misoriented, further supporting the idea that a cell contact-dependent mechanism controls the AB division axis.

Next, we depleted chiral cortical flow. Previous studies increased or decreased chiral cortical flow by manipulating RhoA activity through weak knockdown of a RhoGEF or RhoGAP (Pimpale et al., 2020). However, altering RhoA signaling may have confounding effects on other pathways. A recent study showed that CYK-1/formin is required for chiral cortical flow in AB lineage cells (Middelkoop et al., 2021). We therefore depleted chiral cortical flow using *cyk-1(RNAi)*, rather than directly manipulating RhoA activity. As expected, chiral cortical flow was nearly abolished in *cyk-1(RNAi)* embryos (Figure 2B and Figure S2). Notably, this knockdown condition did not prevent cell division. We found that loss of chiral cortical flow did not affect the initial AB division axis (Figure 2C).

Eggshell geometry was manipulated by altering the eggshell aspect ratio, as described previously (Yamamoto and Kimura, 2017). Mutations in *dpy-11* and *lon-1* cause shorter and longer adult body lengths, respectively, and produce low- and high-aspect-ratio eggshells (Figure S3). We found that neither *dpy-11* nor *lon-1* affected the initial AB division axis (Figure 2D). These results suggest that, among the three pathways investigated, the CUL-3–actomyosin-dependent pathway is the sole regulator of the initial AB division axis.

### CUL-3–actomyosin-dependent oriented division and proper eggshell geometry are required for Phase I rotation

Next, we analyzed the roles of the three pathways in Phase I cellular rotation (Figure 3A and 3B). We found that *cul-3(RNAi)* embryos exhibited a high penetrance of Phase I rotation defects. Although a previous study suggested that chiral cortical flow positively regulates the angular velocity of AB cell rotation (Pimpale et al., 2020), we observed a slight increase in total Phase I rotation in *cyk-1(RNAi)* embryos. Furthermore, we found that *dpy-11* mutants exhibited defective Phase I rotation, whereas no detectable defects were observed in *lon-1* mutants. The extent of Phase I rotation strongly correlated with the initial division-axis orientation at early anaphase in *cul-3(RNAi)* embryos, but not in *dpy-11* mutants (Figure 3C). In contrast, the extent of Phase I rotation correlated with the eggshell aspect ratio in *dpy-11* mutants (Figure 3D). Although *dpy-11* mutants with eggshell aspect ratios within the normal range did not exhibit Phase I rotation defects, those with aspect ratios less than 1.5 showed defective Phase I rotation (Figure 3E). These results suggest that Phase I rotation requires CUL-3–actomyosin-dependent oriented division and elongated eggshell geometry, whereas chiral cortical flow is dispensable.

**Figure 3.**
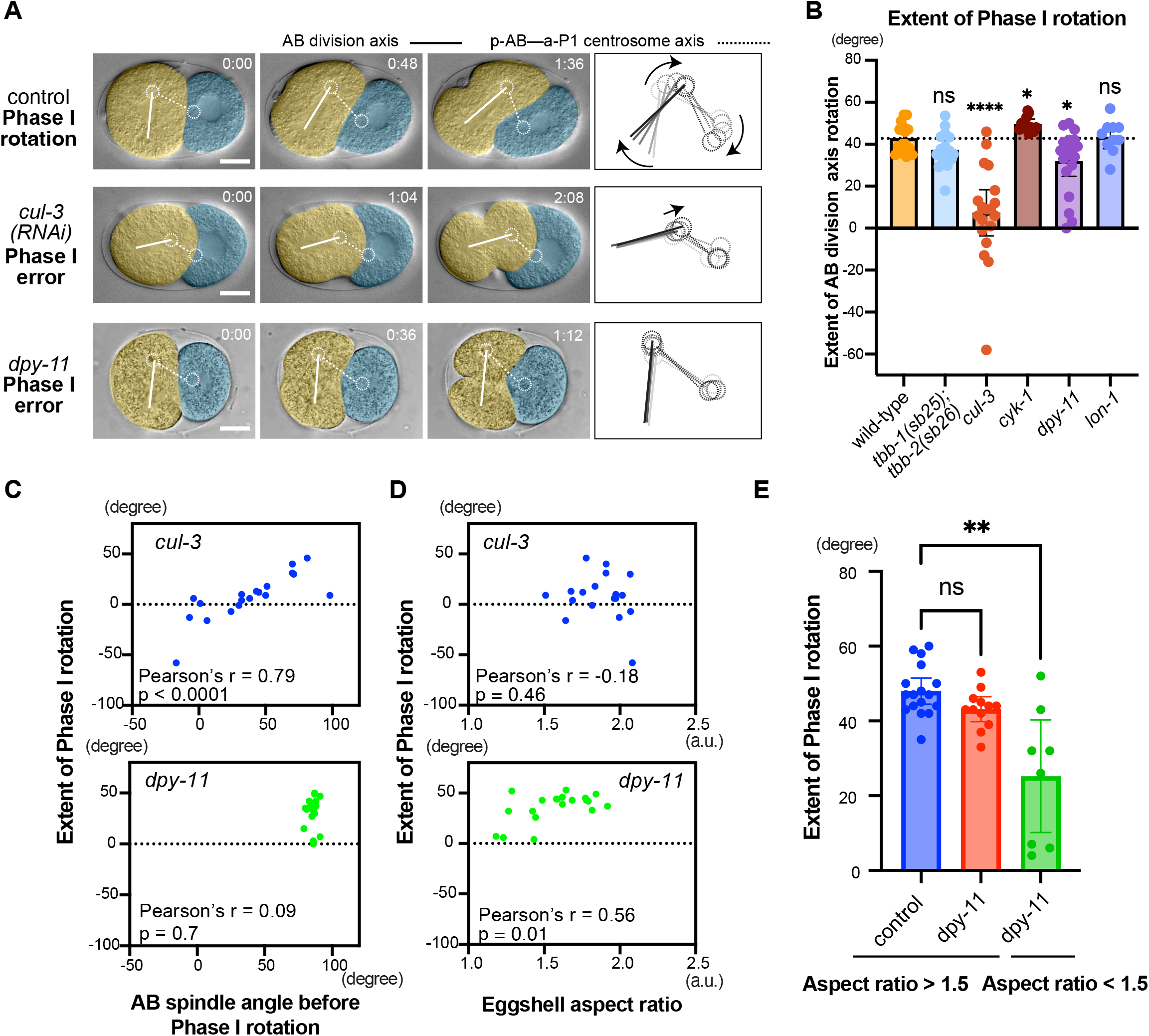
CUL-3–actomyosin-dependent oriented division and elongated eggshell geometry are required for Phase I rotation. **(A**) Cellular rotation during early cytokinesis, corresponding to the Phase I rotation period. Only the posterior AB and anterior P_1_ centrosomes are marked with dotted circles. Solid and dotted lines indicate the AB division axis and an indicator of the dorsal–ventral axis, respectively. Scale bars, 10 µm. Times are relative to the onset of AB cell elongation during anaphase. (**B**) Extent of Phase I AB cell rotation measured by division-axis tilt. (**C**-**D**) Correlation between extent of Phase I rotation and AB spindle angle at early anaphase (before Phase I rotation) (C) and eggshell aspect ratio (D). (**E**) Extent of Phase I rotation in embryos with eggshell aspect ratios greater than or less than 1.5. ****, **, *, and ns indicate p < 0.0001, p < 0.01, p < 0.05, and p > 0.05, respectively, using Brown–Forsythe and Welch ANOVA with Dunnett’s T3 test in (B) and Kruskal-Wallis test with Dunn’s multiple comparison test in (D).

### CUL-3–actomyosin-dependent oriented division is required for Phase II rotation

We also tested the roles of the three pathways in Phase II rotation (Figure 4). In control embryos, the AB division axis rotated during P_1_ cytokinesis (Figure 4A). As suggested by the *div-1*(RNAi) analysis in Figure S1, Phase II rotation is closely linked to P_1_ cytokinesis. When we defined the dorsal and ventral AB–P_1_ boundaries relative to the position of the AB cleavage furrow, we observed that the dorsal AB–P_1_ boundary shifted posteriorly as the P_1_ cleavage furrow invaginated, whereas the ventral AB–P_1_ boundary shifted anteriorly as P_1_ elongated during cytokinesis (Figures 1C and 4B).

**Figure 4.**
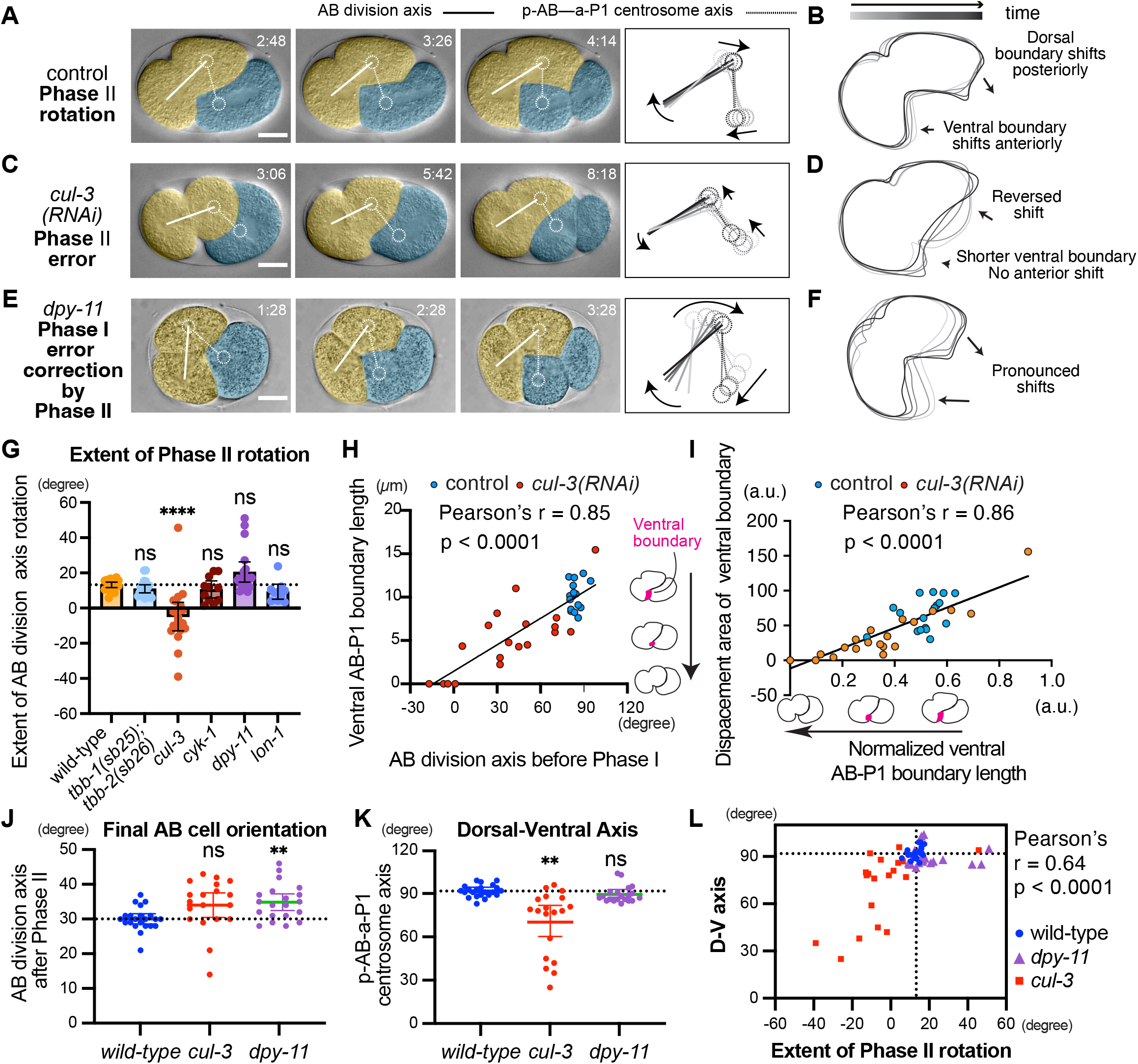
CUL-3–actomyosin-dependent oriented division is required for Phase II rotation. (**A**, **C**, **E**) Cellular rotation during P_1_ cytokinesis, corresponding to the Phase II rotation period. Only the posterior AB and anterior P_1_ centrosomes are marked with dotted circles. Solid and dotted lines indicate the AB division axis and an indicator of the dorsal–ventral axis, respectively. Scale bars, 10 µm. Times are relative to the onset of AB cell elongation during anaphase. (**B**, **D**, **F**) Trajectories of AB cell outlines during Phase II rotation. (**G**) Extent of Phase II AB cell rotation measured by division-axis tilt. (**H**) Relationship between the AB division axis immediately before Phase I rotation and ventral AB–P_1_ boundary length. (**I**) Relationship between normalized ventral AB–P_1_ boundary length (relative to the total boundary length) and the area swept by the ventral boundary. The latter value represents the extent to which the ventral boundary advanced toward the anterior side. (**J**) Final AB cell orientation immediately after Phase II. (**K**) Dorsal-Ventral Axis measured by the posterior AB and anterior P_1_ centrosome axis. (**L**) Relationship between extent of Phase II rotation and the D-V axis. ****, **, *, and ns indicate p < 0.0001, p < 0.01, p < 0.05, and p > 0.05, respectively, by Brown–Forsythe and Welch ANOVA with Dunnett’s T3 test. Dotted lines in graphs indicate wild-type mean.

In contrast, inhibition of CUL-3–actomyosin-dependent oriented division by *cul-3*(RNAi) resulted in defective Phase II rotation (Figures 4C and 4G), whereas defects in chiral cortical flow or eggshell geometry did not affect the extent of Phase II rotation (Figure 4F). In *cul-3(RNAi)* embryos, both the posterior centrosome in AB and the anterior centrosome in P_1_ shifted anteriorly during P_1_ cytokinesis, unlike in control embryos (Figure 4C). Consistently, P_1_ cytokinesis coincided with an anterior shift of the dorsal AB–P_1_ boundary and limited displacement of the ventral AB–P_1_ boundary (Figure 4D). We found that the ventral AB–P_1_ boundary was shorter in *cul-3*(RNAi) embryos than in controls (control, 10.1 ± 1.62 µm; *cul-3*(RNAi), 5.14 ± 4.00 µm; p < 0.0001, Welch’s *t*-test). Moreover, the ventral AB–P_1_ boundary became shorter as the initial AB division-axis defect became more severe (Figure 4H; Pearson’s correlation coefficient, r = 0.85; p < 0.0001; n = 35). This is a geometrically expected outcome, as an A–P-oriented AB division axis positions the anterior part of the dividing AB cell away from the AB–P_1_ boundary (see Figure 3A, second row, and Figures 4A and 4C).

These observations further support the idea that P_1_ cytokinesis induces Phase II rotation by pushing against the ventral AB–P1 boundary. Consistently, reduced ventral AB–P_1_ boundary length correlated with reduced movement of the ventral boundary during P_1_ cytokinesis (Figure 4I; Pearson’s correlation coefficient, r = 0.86; p < 0.0001; n = 35). Together, these results suggest that P_1_ cytokinesis-dependent pushing against the ventral AB–P_1_ boundary drives Phase II rotation and that this process is defective in *cul-3*(RNAi) embryos.

### Phase I defects become fatal D-V axis errors when Phase II rotation is defective

As defects in CUL-3–actomyosin-dependent oriented division cause both A–P-orthogonal division-axis defects and Phase I rotation defects, it remained possible that Phase I rotation defects themselves lead to defective Phase II rotation. However, Phase I rotation errors in *dpy-11* mutants were completely rescued during Phase II rotation (Figures 4E and 4G). Although not statistically significant, Phase I errors in *dpy-11* mutants were often compensated for by an increased extent of Phase II rotation (Figure 4F). These results suggest that Phase I errors per se are not necessarily fatal to dorsal– ventral axis establishment, as they can be corrected by Phase II rotation. They further suggest that Phase I rotation errors persist in *cul-3(RNAi)* embryos due to defective Phase II rotation.

Is Phase II rotations indeed necessary for D-V axis establishment? The distinct trajectories of AB cell orientation in wild-type, *dpy-11*, and *cul-3(RNAi)* provide some insight. In wild-type embryos, the initial division axis is A-P-orthogonal, and the AB cell subsequently undergoes normal Phase I and Phase II rotations. In *dpy-11*, the initial division axis is normal, and Phase I errors are corrected during Phase II. In *cul-3(RNAi)*, the initial division axis is more horizontal, and both Phase I and Phase II rotations are defective. Paradoxically, these distinct trajectories lead to similar final AB cell orientations after Phase II in all three conditions (Figure 4J). However, only *cul-3(RNAi)* exhibited defective D-V axis orientation, measured by the line connecting the posterior centrosome in AB and the anterior centrosome in P_1_ (“the p-AB-a-P1 centrosome axis” in Figure 4C; Figure 4K). We found that reduced or reversed Phase II rotation is associated with defective D-V axis orientation (Figure 4L; Pearson’s r = 0.64, p < 0.0001, n = 59). Indeed, microscopy images show that the p-AB-a-P1 centrosome axis, the future D-V axis, is dynamically reoriented during Phase II rotation and reaches approximately 90° at the end of Phase II in both control and *dpy-11* embryos (Figure 4A and 4E, dotted lines). On the other hand, due to incorrect pushing of the dorsal AB-P_1_ boundary during P_1_ division, the AB cell undergoes reversed Phase II rotation, leading to defective reorientation of the p-AB-a-P1 centrosome axis (Figure 4C, dotted lines). These results suggest that it is the cellular rotation process, rather than the final AB cell orientation, that ensures correct D-V axis establishment.

### CUL-3-actomyosin-dependent oriented division corrects division axis before and during cytokinesis but is not sufficient to induce Phase I rotation

Are Phase I and Phase II rotation defects in *cul-3(RNAi)* embryos entirely attributed to the initial misorientation of the division axis? We previously demonstrated that wild-type AB cells possess a mechanism for correcting division-axis orientation errors, as the attachment of an adhesive rhodamine-coated bead induces oriented division of isolated AB cells parallel to the cell–bead boundary (Figure 5A) (Sugioka and Bowerman, 2018). Rhodamine-coated beads adhere to cells through electrostatic interactions and are hereafter referred to as adhesive beads. Even when isolated AB cells were brought into contact with adhesive beads at random positions, this mechanism corrected inappropriate division axes during cytokinesis (Figure 5B). Although *cul-3(RNAi)* embryos exhibited division-axis orientation defects before cytokinesis, their ability to correct these defects during cytokinesis was also impaired (Figure 3A). Thus, CUL-3–actomyosin-dependent oriented division likely operates both before and during cytokinesis in vivo. These findings suggest that Phase I and Phase II rotation defects in *cul-3(RNAi)* embryos cannot be attributed solely to the initial misorientation of the division axis.

**Figure 5.**
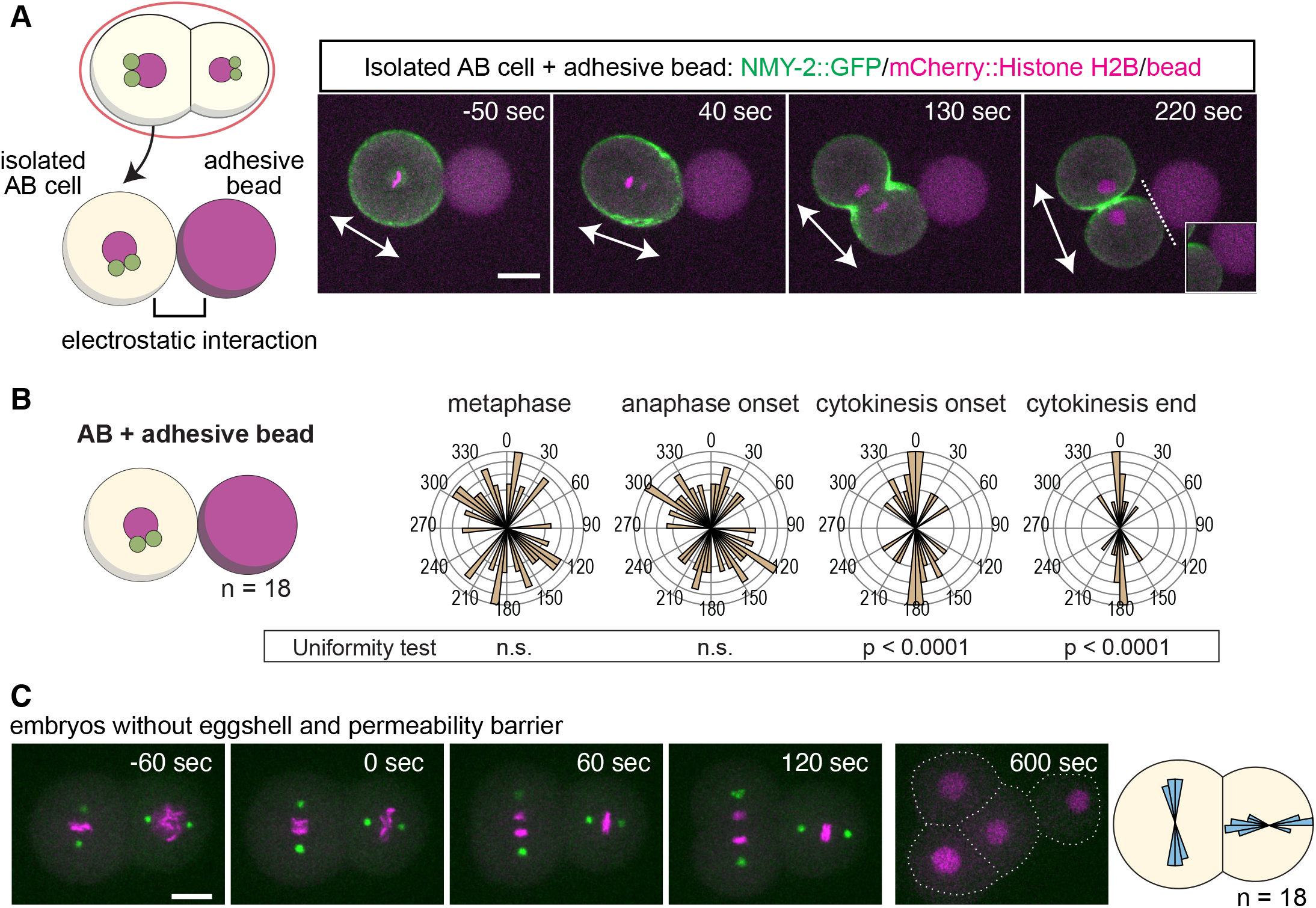
CUL-3–actomyosin-dependent oriented division can correct the AB division axis during cytokinesis but is not sufficient to induce Phase I or Phase II rotation. (**A**) Contact with adhesive beads orients the AB division axis. AB cells were isolated at the two-cell stage and brought into contact with adhesive beads. NMY-2::GFP, myosin II; mCherry::Histone H2B, chromosomes; rhodamine, beads. Arrows indicate the AB division axis. Dotted lines indicate cell–bead contacts. (**B**) AB division-axis orientation from metaphase until the end of cytokinesis. P values are based on Rayleigh’s test, where p < 0.05 rejects the null hypothesis that the angle distribution is uniform. (**C**) AB division-axis orientation in embryos after removal of the eggshell and permeability barrier. Time is relative to anaphase onset. The right panel shows spindle orientation at AB anaphase onset and P_1_ metaphase. Scale bar, 10 µm.

In the in vitro adhesive bead assay, the AB division axis remained parallel to the cell–bead boundary once it had achieved the correct orientation and did not undergo further rotation (Figure 5B). Furthermore, removal of the eggshell and permeability barrier led to defective Phase I and Phase II rotations (Figure 5C), consequently resulting in a T-shaped four-cell-stage arrangement (Figure 6B). Thus, the CUL-3-actomyosin-dependent oriented division mainly regulates the AB division axis before and during cytokinesis, but does not induce Phase I rotation per se. As division-axis correction occurs only after anaphase onset in the in vitro assay (Figure 5B), distinct mechanisms may regulate AB division-axis orientation before and during cytokinesis. Thus, we refer to the latter mechanism as the contact-dependent division-axis correction mechanism.

**Figure 6.**
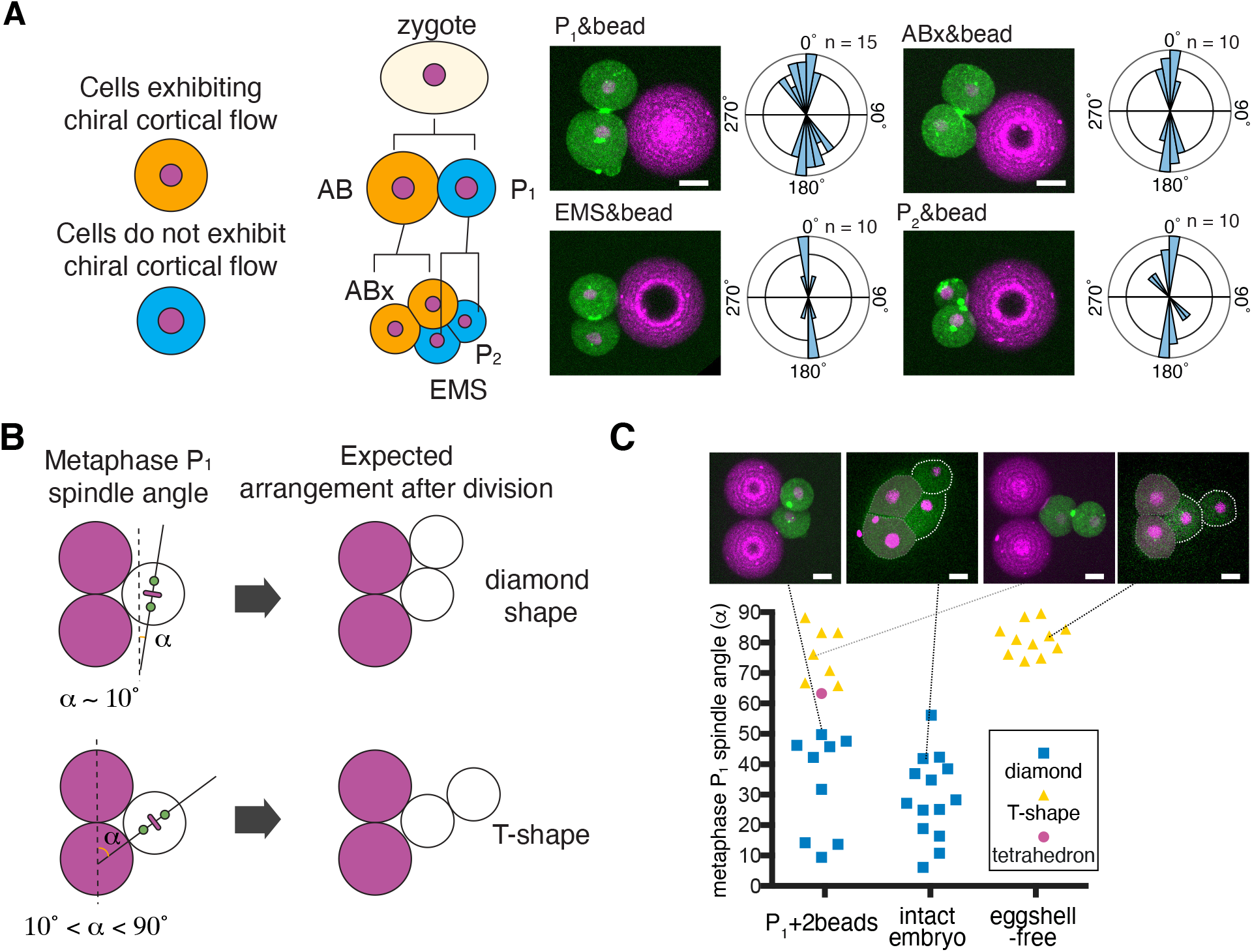
The contact-dependent division-axis correction mechanism regulates the four-cell-stage arrangement. (**A**) Lineage-wide analysis of the contact-dependent division-axis correction mechanism. Isolated embryonic founder cells were attached to adhesive beads at random positions before metaphase, and division-axis orientation was followed through cytokinesis. Angular plots indicate division-axis orientation at the end of cytokinesis. (**B**) Experimental scheme simulating P_1_ division after Phase I rotation. An isolated P_1_ cell was attached to two adhesive beads. The angle of the metaphase spindle relative to the line passing through the two beads is denoted as α. Assuming linear division without division-axis correction, a diamond-shaped arrangement after P_1_ division is possible only when α is minimal. (**C**) Distribution of P_1_ spindle orientation and the resulting cell–bead arrangement. Scale bars, 10 µm.

### The contact-dependent division-axis correction mechanism operates in all embryonic founder cells during cytokinesis

Cortical and spindle regulation are distinctly controlled in different cellular lineages. For example, chiral cortical flow is observed only in the anterior lineage cells, including AB and its descendants (Figure 6A, schematic) (Pimpale et al., 2020). We therefore asked whether the contact-dependent division-axis correction mechanism is restricted to the AB lineage. To this end, we isolated AB daughter cells, denoted as ABx because ABa and ABp cannot be distinguished after isolation, as well as P_1_ and the P_1_ daughter cells EMS and P2, and attached them to adhesive beads (Figure 6A). Surprisingly, in all cases, the division axis became parallel to the cell–bead contact by the end of cytokinesis. These results suggest that the contact-dependent division-axis correction mechanism operates in all embryonic founder cells during cytokinesis.

Division-axis correction during P_1_ cytokinesis is of particular interest because it may contribute to the four-cell-stage blastomere arrangement. To test this, we isolated P_1_ cells and attached them to two adjacent adhesive beads, thereby simulating the arrangement after Phase I rotation. We quantified metaphase spindle orientation by measuring the angle between the mitotic spindle and the line passing through the two beads, denoted as α. To establish a diamond-shaped arrangement after P_1_ division, in which all five contacts are formed among cells and beads, the initial P_1_ spindle must be nearly parallel to the cell–bead contact site from the beginning (Figure 6B, top). Any deviation of α beyond this range inevitably leads to formation of a T-shaped arrangement, in which only four contacts are formed because one daughter cell moves away from the adhesive beads, resulting in catastrophic failure of dorsal–ventral axis establishment (Figure 6B, bottom). However, in intact embryos, we often observed P_1_ metaphase spindle angles as large as 50°, yet these embryos still formed a diamond-shaped arrangement (Figure 6C, intact embryos).

Experimentally, we quantified metaphase spindle orientation and then followed the arrangement of the bead–cell aggregate after cytokinesis. Strikingly, diamond-shaped arrangements formed even when α was as large as 50° (Figure 6C, P_1_+2beads). When α exceeded 50°, division-axis errors were no longer fully corrected, resulting in T-shaped arrangements similar to those observed in embryos lacking the eggshell and permeability barrier (Figure 6C). These in vitro experiments suggest that the division-axis correction mechanism during P1 cytokinesis may influence cell arrangement at the four-cell stage, which is critical for dorsal–ventral axis establishment. Our data also suggest that complete suppression of the T-shaped arrangement requires eggshell confinement.

### Ras–MAPK signaling is a novel regulator of Phase I rotation

Our recent RNAi screen identified the Ras–MAPK signaling adaptor *sem-5*/Grb2 as being required for AB cell rotation (Gough et al., 2025). Ras–MAPK signaling interacts with RhoA signaling, a master regulator of cytokinesis, in the context of vulval cell-fate specification (Canevascini et al., 2005) (Figure 7A). We therefore tested during which phase of dorsal–ventral axis establishment *sem-5* and other Ras–MAPK signaling components function.

**Figure 7.**
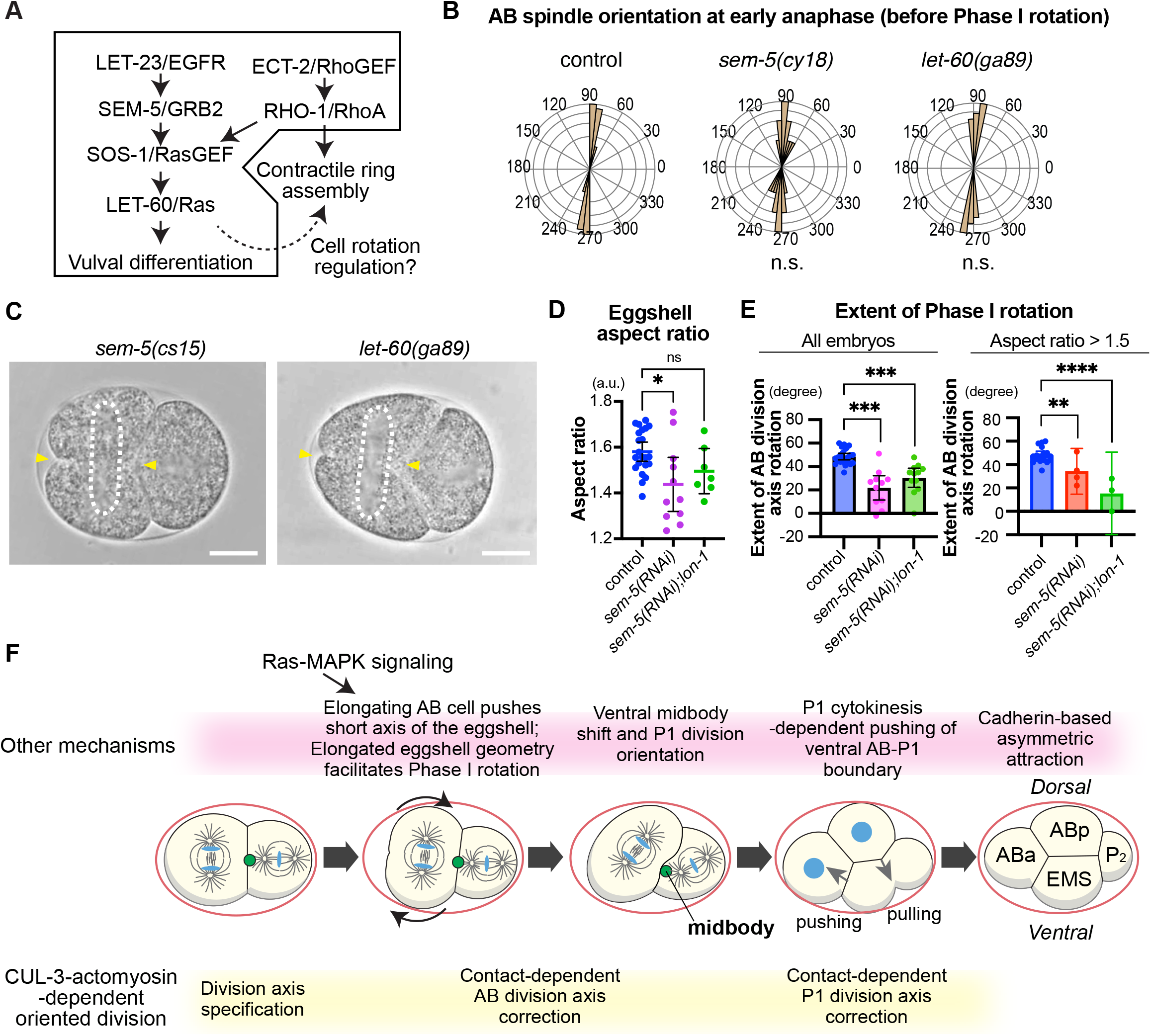
Ras-MAPK signaling is involved in Phase I rotation. **(A**) Genetic interaction between Ras–MAPK signaling and RhoA signaling. The known interaction between these pathways during vulval cell-fate specification is shown in the black box. (**B**) A–P-axis-orthogonal division-axis orientation was not affected in the reduction-of-function mutant sem-5(cs15) or the gain-of-function mutant let-60(ga89). Control data are identical to those shown in Figure 2C. P values were calculated using the Mardia–Watson–Wheeler test for equal distributions. (**C**) Defects in Phase I AB cell rotation in *sem-5(cs15)* and *let-60(ga89)* embryos. Scale bar, 10 µm. (**D**) Eggshell aspect ratio in *sem-5(RNAi)* embryos. (**E**) Extent of Phase I AB cell rotation measured by division-axis tilt. Only embryos with an eggshell aspect ratio greater than 1.5 were included in the right panel. ***, *, and ns indicate p < 0.001, p < 0.05, and p > 0.05, respectively, by Brown–Forsythe and Welch ANOVA with Dunnett’s T3 test. (**F**) Proposed mechanism of cellular rotation during dorsal–ventral axis establishment; see text for details.

We found that neither the reduction-of-function *sem-5* mutant *sem-5(cy15)* nor the gain-of-function Ras mutant *let-60(ga89)* exhibited detectable defects in specification of the A–P-orthogonal division axis in AB (Figure 7B). However, both mutants exhibited defects in Phase I rotation (Figure 7C). Ras–MAPK signaling could regulate Phase I rotation through effects on eggshell geometry. Consistent with this possibility, we found that *sem-5*(RNAi) embryos exhibited a reduced eggshell aspect ratio. To test whether the Phase I rotation defects were due to altered eggshell geometry, we used the *lon-1* mutant, which has a larger eggshell aspect ratio, as shown in Figure S3. As expected, the eggshell aspect ratio was restored to normal levels in *sem-5(RNAi); lon-1* embryos (Figure 7D). However, Phase I rotation remained defective in these embryos (Figure 7E, left). Furthermore, both *sem-5(RNAi)* and *sem-5(RNAi); lon-1* embryos with eggshell aspect ratios greater than 1.5, within the wild-type range, still exhibited Phase I rotation defects (Figure 7E, right). These results suggest that Ras–MAPK signaling specifically regulates Phase I rotation rather than acting indirectly through eggshell geometry (Figure 7E).

## Discussion

In this study, we analyzed the mechanism underlying cellular rotation during dorsal-ventral axis establishment in *C. elegans*. We identified two phases of cellular rotation: the first occurs during AB cytokinesis, whereas the second occurs during P_1_ cytokinesis. We found that Phase I rotation is regulated by CUL-3–actomyosin-dependent oriented division and eggshell geometry. Phase I errors can be fully corrected by subsequent Phase II rotation, underscoring the role of two-step cellular rotation as a robust fail-safe mechanism. However, defective CUL-3-actomyosin-dependent oriented division compromises this error correction mechanism during Phase II, leading to fatal D-V axis misorientation. Adhesive bead experiments further showed that CUL-3– actomyosin-dependent oriented division operates both before and during cytokinesis, with the latter mechanism functioning in all embryonic founder cells. Additional bead experiments suggested that the division-axis correction mechanism in P_1_ facilitates the correct four-cell-stage arrangement. Finally, we found that Ras–MAPK signaling is involved in Phase I rotation. Taken together, this study defines distinct steps of cellular rotation and identifies the mechanisms involved in each step.

We propose the following mechanism of D-V axis establishment (Figure 7F). First, CUL-3-actmyosin-dependent oriented division regulates A-P-orthogonal division axis. As the cell elongates and hit the eggshell at the right angle, AB cell undergoes Phase I rotation. The elongated geometry of the eggshell facilitates Phase I rotation, whereas a more spherical eggshell, as observed in *dpy-11* mutants, may impair Phase I rotation by reducing the geometric preference for a particular orientation. Ras-MAPK signaling and RhoA signaling may regulate AB cell mechanical property such as eggshell pushing force. Contact-dependent division axis correction mechanism should operate during AB cytokinesis to maintain the correct division axis horizontal to the cell-cell contact site. As this mechanism only ensures that the AB division axis is oriented parallel to the cell–cell contact interface, it would not prevent cellular rotation. Once AB cell rotates, midbody shifts ventrally and orients P_1_ spindle. As P_1_ undergoes cytokinesis, P_1_ cell pushes ventral AB-P_1_ boundary and regulates Phase II rotation. P_1_ contractile ring might also pull the dorsal side of AB blastomere. Contact-dependent division axis correction mechanism also help ensuring the diamond shaped 4-cell stage arrangement and should work with a previously described cell adhesion-dependent asymmetric attraction mechanism (Yamamoto and Kimura, 2017).

Our study demonstrates that chiral cortical flow is not a universal driver of cellular rotation and that distinct mechanisms of cellular rotation may cooperate during embryogenesis. A pioneering study demonstrated the role of chiral cortical flow in *C. elegans* embryogenesis at the 4-cell stage (Naganathan et al., 2014). At this stage, the daughter cells of AB, ABa and ABp, undergo clockwise tilting of their division axes when viewed from the dorsal side, which determines organismal left–right handedness (Wood, 1991). Increasing or decreasing cortical flow chirality through RNAi-mediated knockdown of a RhoA inhibitor or activator, respectively, resulted in corresponding changes in the rate of division-axis skewing (Naganathan et al., 2014). Loss of chiral cortical flow in *cyk-1(RNAi)* embryos also resulted in the loss of division-axis skewing (Middelkoop et al., 2021). Similar experiments were conducted in AB cells at the two-cell stage, where increases and decreases in chiral cortical flow correlated with corresponding changes in the rate of AB division-axis skewing (Pimpale et al., 2020). However, our previous study did not identify a role for chiral cortical flow in AB cell rotation (Sugioka and Bowerman, 2018). We reported that cortical flow from the polar region toward the cleavage furrow (defined as x-axis flow, or contractile ring-directed flow) and flow along the perpendicular axis (y-axis flow) exhibited distinct asymmetries during AB cell division. The y-axis flow asymmetry corresponds to chiral cortical flow. We found that *cul-3(RNAi)* abolished the asymmetry in x-axis flow as well as AB cell rotation without impairing chiral cortical flow. These results suggested that chiral cortical flow was not required for AB cell rotation (Sugioka and Bowerman, 2018). Consistent with this finding, the present study demonstrates that the loss of chiral cortical flow in *cyk-1(RNAi)* embryos does not affect the final orientation of the AB division axis, raising the possibility that distinct mechanisms of cellular rotation operate in AB cells at the 2-cell stage and in ABa and ABp cells at the 4-cell stage. Extending this observation, we speculate that the relative contributions of these mechanisms may vary depending on cell-type-specific requirements during embryogenesis, rather than being governed by a single, unified mechanism.

This study has several limitations. First, embryos were imaged on agarose pads, where they are slightly compressed. Compression is known to alter cortical actomyosin dynamics (Singh et al., 2019), and the relative contributions of different pathways may differ slightly between compressed and compression-free conditions. We also did not investigate the contributions of other known mechanisms, including the dynein-dependent centrosome separation pathway in AB (Bondaz et al., 2019), the dynein-dependent mechanism that enables long-axis division in AB (Middelkoop et al., 2024), and compression-induced rotational cortical flow that biases the midbody toward the ventral side (Singh and Pohl, 2014). We excluded the former two mechanisms because published results suggest that they are unlikely to affect the outcome of dorsal–ventral axis establishment. We also did not examine the mechanism underlying midbody displacement, which allowed us to focus specifically on the AB cell rotation process.

This study uncovered the significance of cellular rotation in dorsal–ventral axis establishment and identified new roles for pathways that regulate this process. However, the precise physical and molecular mechanisms underlying these events remain unclear. Future studies should elucidate the mechanistic roles of these pathways in greater detail.

## Materials and Methods

### Maintenance of worm strains

All *C. elegans* strains were cultured and maintained using standard methods (Brenner, 1974). The following knock-in or transgenes were used: *nmy-2(cp13*[*nmy-2*::GFP + LoxP]) (knock-in reporter, (Dickinson et al. 2013)), *cpIs56*[*mex-5*p::TagRFP-T::PLC(delta)-PH::*tbb-2* 3’UTR + *unc-119* (+)] (transgene, (Heppert et al., 11 07, 2016)), *sas-7*(*or1940*[GFP::*sas-7*]) (knock-in, (Sugioka et al., 2017)), and ltIs37[pie-1p::mCherry::his-58 + unc-119(+)] (transgene, (McNally et al., 2006)).

The following homozygous mutant strains were analyzed: *sem-5(cy15)*, *let-60(ga89)*, *dpy-11(e207)*, and *lon-1(e185)*. The temperature-sensitive *let-60(ga89)* mutant was maintained at 15°C and shifted to the restrictive temperature of 25°C overnight before imaging analysis.

### RNAi

Feeding RNAi was performed for control, *cul-3*, *cyk-1*, *div-1*, *hmr-1*, and *sem-5* at 25°C using standard methods (Ahringer, 2006). dsRNA expression in the *Escherichia coli* strain HT115 was induced by overnight incubation at 37°C on Nematode Growth Medium (NGM) plates containing 10 µg/mL ampicillin and 1 mM isopropyl β-D-1-thiogalactopyranoside (IPTG). For control RNAi, a bacterial strain carrying the empty vector L4440 was used.

For control, *cul-3*, *hmr-1*, and *sem-5* RNAi, gravid adults were bleached with hypochlorite solution, and the resulting purified embryos were incubated in M9 buffer overnight to obtain developmentally arrested L1-stage worms. These worms were grown on feeding RNAi plates, and their embryos were analyzed. For *cul-3* RNAi, L1 larvae were grown on feeding RNAi plates, and sterile F1 adults were crossed with *cul-3*(RNAi) males to obtain embryos for analysis. For *cyk-1* and *div-1* RNAi, L4 worms were incubated on RNAi plates overnight before imaging.

### Live-imaging

To obtain embryos for live imaging by brightfield microscopy, gravid adults were dissected on a coverslip in a 5-µL droplet of M9 buffer. The coverslip was then gently placed onto a 2% agarose pad mounted on a glass slide, and the edges were sealed with petroleum jelly (Vaseline) to prevent dehydration. Live imaging was performed at room temperature (∼22.5°C) using a brightfield compound microscope (B290-TB; OPTIKA) equipped with a 100× oil-immersion objective lens (NA 1.2) and controlled by OPTIKA Vision Lite software. Images were acquired every 2 s beginning at the early two-cell stage. For Figures 1, S1, 2B, and S2, dissected embryos were mounted on 2% agarose pads and imaged using an Olympus IX83 microscope (Olympus/Evident) equipped with a CSU-W1 spinning-disk confocal unit (Yokogawa), a Prime95B scientific CMOS camera (Photometrics), a NANO-Z piezoelectric stage (Mad City Labs), and a UPLSAPO60XS2 silicon-immersion objective lens (60×, NA 1.3; Olympus/Evident), all controlled by CellSens Dimension software (Olympus/Evident). Silicone immersion oil (Z81114; refractive index, 1.406 at 23°C; Olympus/Evident) was used as the immersion medium. Imaging was performed using 488-nm and 561-nm diode-pumped lasers with a 150-ms exposure time, 1-µm z-step size, 28 slices per frame, and a 5-s frame interval.

For Figures 5 and 6, eggshell-free embryos and isolated blastomeres were placed on an uncoated coverslip in a droplet of Shelton’s growth medium. Imaging was performed using a CSU-W confocal unit with Borealis illumination (Andor) and an iXon Ultra 897 EMCCD camera (Andor) mounted on a Leica DMi8 inverted microscope (Leica). NMY-2::GFP and mCherry::Histone H2B signals were acquired every 15 s with 2-µm z-spacing.

#### Eggshell removal, blastomere isolation, and adhesive bead assays

Eggshell and permeability barrier removal was performed as previously described (Edgar and Goldstein, 2012) with some modifications(Hsu et al., 2019). Gravid adults were cut open in egg salt buffer, and the released embryos were incubated in hypochlorite solution [75% Clorox (Clorox) and 2.5 N KOH] for 50 s. After two washes with Shelton’s growth medium, embryos were devitellinized by mouth pipetting using hand-drawn microcapillary tubes (10 µL; Kimble Glass Inc.).

Carboxyl-modified polystyrene beads of 30-µm diameter (10 mg; KISKER BIOTECH GmbH & Co.) were washed twice with 100 mM 2-(N-morpholino)ethanesulfonic acid (MES) buffer, pH 6.5, and incubated in 1 mL MES buffer containing 10 mg 1-ethyl-3-(3-dimethylaminopropyl)carbodiimide (EDAC) for 15 min at room temperature, as described previously (Hsu et al., 2019). The beads were washed twice with phosphate-buffered saline (PBS) and incubated in 0.5 mL PBS containing 0.05 µg Rhodamine Red-X succinimidyl ester (Thermo Fisher Scientific) for 5 min. The beads were then washed twice and stored in PBS at 4°C before use.

Cells isolated before prometaphase were attached to either one or two rhodamine-coated adhesive beads before imaging. For one-cell/two-bead assays, two beads were first brought together and then attached to the isolated cell. All manipulations were performed using a mouth pipette as described previously (Hsu et al., 2019).

### Image analysis

To generate images of spindle pole trajectories in Figures 1B, S1A, and 3A, centrosome positions were manually marked, and multiple frames were superimposed. Similarly, to generate images of cell outline trajectories in Figures 4B, 4D, and 4F, AB cell outlines were manually marked, and multiple frames were superimposed.

The area swept by the ventral AB–P1 boundary was measured by identifying the area through which the AB cell shifted during Phase II relative to the onset of Phase II.

The dorsal-ventral axis is measured by the line connecting the posterior centrosome in AB and the anterior centrosome in P_1_ immediately after Phase II rotation.

For cortical flow analysis, images were flattened by maximum-intensity projection of the 10 surface z-slices closest to the objective lens. Cortical flow was estimated using PIVlab software (Thielicke and Sonntag, 2021). The average velocity of cortical foci along the vertical axis was calculated to quantify cortical flow along the axis aligned with the cleavage furrow. Velocities on the left and right sides relative to the cleavage furrow were defined as *v*y,left and *v*y,right, respectively. Chiral cortical flow velocity, representing the degree of counter-rotating flow, was then measured by subtracting *v*y,right from *v*y,left.

For the division axis analysis in Figure 5, 3D projections of time-lapse images were rotated to identify a viewing angle at which the cells and beads did not overlap. The cell-bead contact sites were then manually annotated. The division axis was determined based on the orientation of chromosome markers, using either the metaphase plate or segregating chromosomes. In Figure 5B, the division axis is shown relative to the cell-bead contact site.

### Statistics

Statistical analyses were performed using Prism 9 (GraphPad Software). The resulting p values were corrected for multiple comparisons. Symbols are defined as follows: ****, p < 0.0001; ***, p < 0.001; **, p < 0.01; *, p < 0.05; and ns, p > 0.05. Center values in graphs represent means. Sample sizes were not predetermined by statistical analysis. No blinding was performed for any analysis.

## Data and resource availability

All relevant data and details of resources can be found within the article and its supplementary information.

## Acknowledgements

We thank Caenorhabditis Genetics Center (funded by the NIH Office of Research Infrastructure Programs; P40 OD010440) for sharing worm strains and the Sugioka lab members for general discussions. This work was supported by the Canadian Institutes of Health Research (Project Grant; PJT-169145), Government of Canada’s New Frontiers in Research Fund (NFRFE-2019-00310), and the Michael Smith Health Research BC (Scholar Award; SCH-2020-0406) to K.S.

