## Supplementary Figures 1-3 for "Combinatorial regulation of cellular rotation during dorsal-ventral axis establishment in *Caenorhabditis elegans*"

### Khor et al., Figure S1

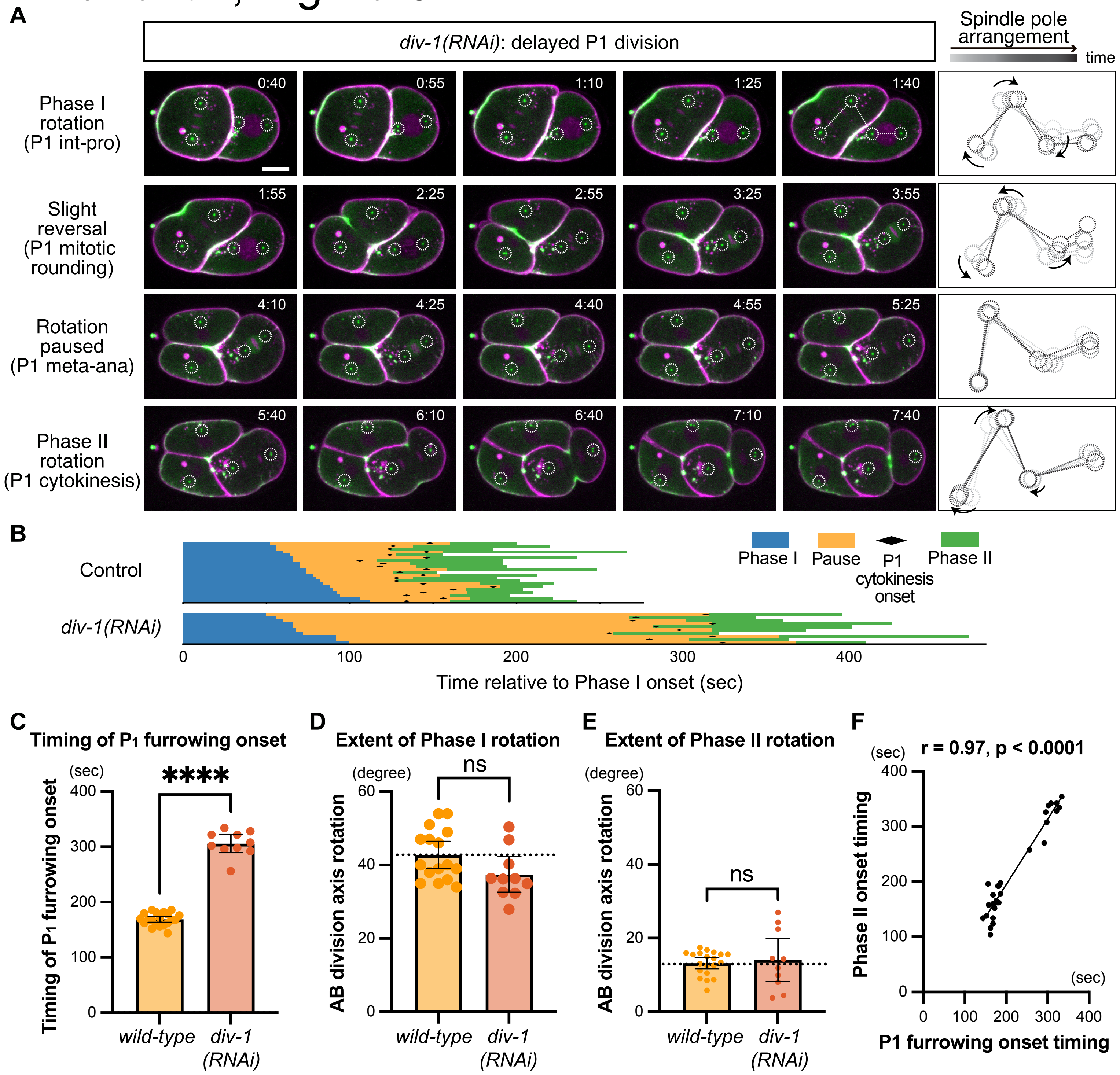


**Figure S1. Phase I rotation occurs independently of P1 division, whereas Phase II rotation coincides with P1 cytokinesis.**

(**A**) Two phases of cellular rotation during AB and P1 divisions in *div-1(RNAi)*. NMY-2 (green, myosin), SAS-7 (green, centriole), TagRFP::PH (magenta, plasma membrane), mCherry::Histone H2B (magenta, chromosome). Dotted circles indicate centrosome positions. Right panels show the trajectories of each centrosome and lines connecting key centrosomes. Times are minutes and seconds relative to anaphase onset. (**B**) Timing of events in individual embryos. Each horizontal bar represents a single embryo, with different colors indicating the duration of each phase. (**C**) Timing of P1 furrowing onset relative to the onset of Phase I rotation. (**D**) Extent of Phase I AB cell rotation. (**E**) Extent of Phase II AB cell rotation. Welch's t-test was used in B-D. (**F**) Correlation between P1 furrowing onset timing and Phase II onset timing. Pearson's correlation coefficient isshown. Scale bars, 10 μm.

### Khor et al., Figure S2

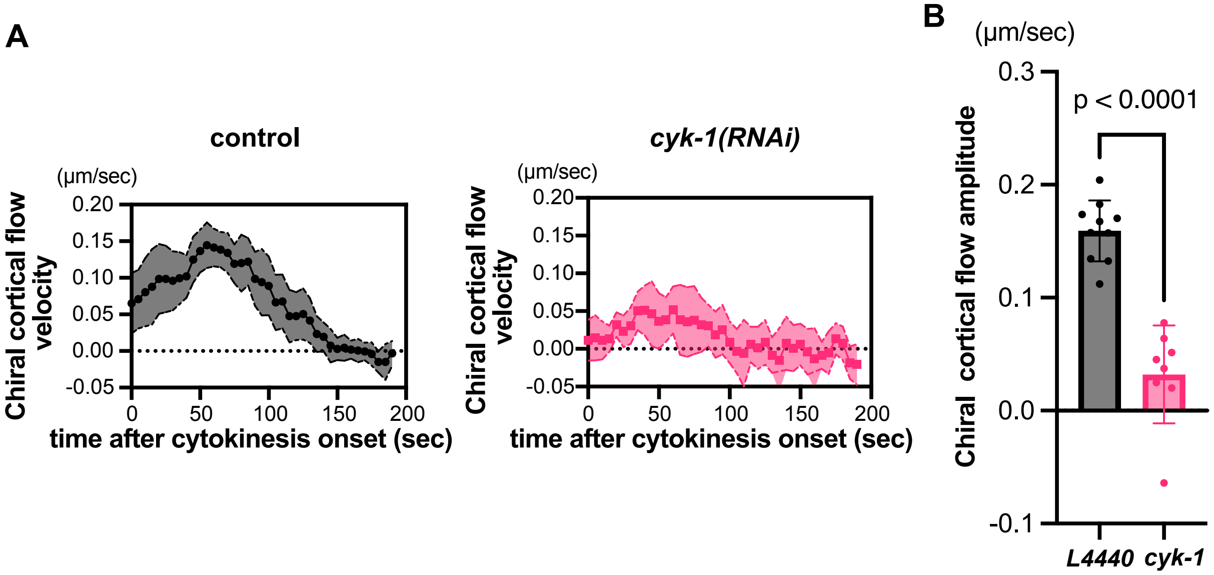


**Figure S2. Depletion of chiral cortical flow by cyk-1(RNAi).**
(A) Quantification of chiral cortical flow velocity, measured as the degree of counter-rotation between the two daughter hemispheres of the AB cell during cytokinesis. (B) Amplitude of chiral cortical flow estimated by Gaussian fitting. P values were calculated using Welch's t-test.

### Khor et al., Figure S3

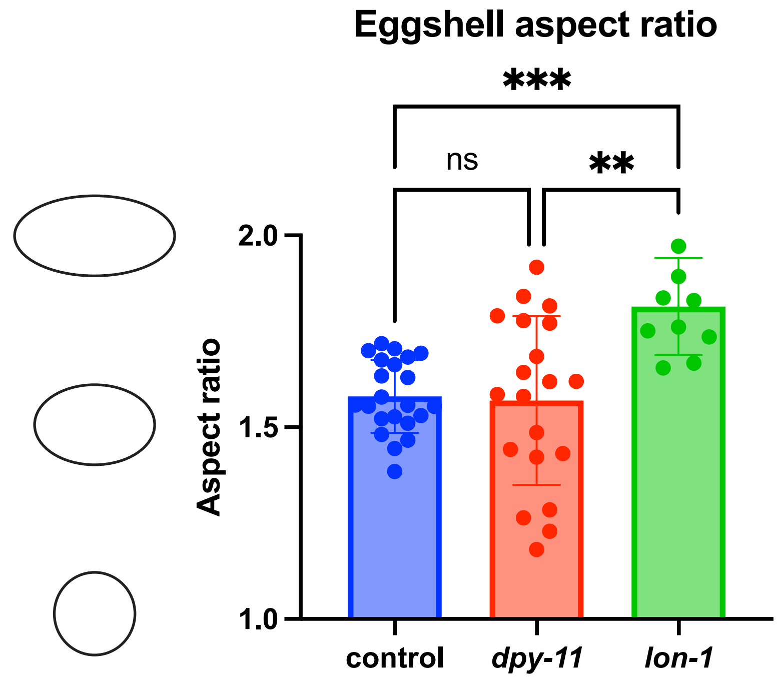


**Figure S3. Genetic manipulation of eggshell aspect ratio.**
Eggshell aspect ratio was measured by dividing the A–P length by the A–P-axis-orthogonal length. dpy-11 mutants exhibited a wider range of aspect ratios compared with control embryos, including a greater proportion of round eggshells. lon-1 mutants exhibited a higher aspect ratio. ***, **, and ns indicate $p < 0.001$, $p < 0.01$, and $p > 0.05$, respectively, by Brown–Forsythe and Welch ANOVA.
